# Alterations in leptin signaling in Amyotrophic Lateral Sclerosis (ALS)

**DOI:** 10.1101/2021.07.14.452319

**Authors:** Agueda Ferrer-Donato, Ana Contreras, Laura M. Frago, Julie A. Chowen, Carmen M. Fernandez-Martos

**Affiliations:** Research Unit of the National Hospital of Paraplegics (UDI-HNP), Toledo, Spain. (A.F.-D.); (CM.F.-M.); Health Research Centre (CEINSA), University of Almeria (UAL), Almeria, Spain. (A.C.); Department of Pediatrics & Pediatric Endocrinology, Hospital Infantil Universitario Niño Jesús, Universidad Autónoma de Madrid; Instituto de Investigación la Princesa. Madrid, Spain; Centro de Investigación Biomédica en Red Fisiopatología de la Obesidad y la Nutrición (CIBEROBN), Instituto de Salud Carlos III, IMDEA Food Institute, CEI UAM + CSIC, Madrid, Spain. (J.A.C.); (L.M.F); Wicking Dementia Research and Education Centre, College of Health and Medicine, University of Tasmania, Hobart, Tasmania, Australia. (CM.F.-M.)

**Keywords:** Neurodegenerative disease, Amyotrophic lateral sclerosis (ALS), Metabolism, Leptin, Long leptin receptor (Ob-Rb), TAR DNA binding protein (TDP-43)

## Abstract

Leptin has been suggested to play a role in amyotrophic lateral sclerosis (ALS), a fatal progressive and neurodegenerative disease. This adipokine has previously been shown to be associated with a lower risk of ALS disease and to confer a survival advantage in ALS patients. However, the role of leptin in the progression of ALS is unknown. Indeed, our understanding of the mechanisms underlying leptin’s effects in the pathogenesis of ALS is very limited, and it is fundamental to determine whether alterations in leptin’s actions take place in this neurodegenerative disease. To characterize the association between leptin signaling and the clinical course of ALS we assessed the mRNA and protein expression profiles of leptin, the long leptin receptor (Ob-Rb) and leptin-related signaling pathways over the time course of the disease (onset and end-stage of disease), in TDP-43^A315T^ mice compared to age-matched WT littermates. In addition, at the selected time-points immunoassay analysis was conducted to characterize plasma levels of total ghrelin, the adipokines (resistin and leptin) and metabolic proteins (plasminogen activator inhibitor type 1 (PAI-1), gastric inhibitory peptide (GIP), glucagon like peptide 1 (GLP-1), insulin and glucagon) in TDP-43^A315T^ mice compared to WT controls. Our results indicate alterations in leptin signaling in the spinal cord and the hypothalamus on the backdrop of TDP-43-induced deficits in mice, providing new evidence about the pathways that could link leptin signaling to ALS.

## INTRODUCTION

Amyotrophic lateral sclerosis (ALS), is an irreversible neurodegenerative disorder characterized by the selective and progressive loss of upper and lower motor neurons of the cerebral cortex, brainstem, and spinal cord (Tapia, 2014). ALS is fast becoming a major health and socioeconomic challenge for many countries across the world; however, no current therapeutic diseasemodifying intervention exists. Although the cellular basis for neurodegeneration in ALS is not yet fully understood, numerous studies have shown that the underlying disease process involves multiple complex genetic and non-genetic factors, including metabolic alterations.

Included amongst the underlying metabolic alterations in ALS is leptin, a polypeptide hormone secreted primarily by adipocytes that exerts an important role in regulating food intake and energy balance through actions in the brain (Stephens et al., 1995; Zhang et al., 1994). Leptin acts by binding to its receptors that are structurally related to the cytokine receptor class I family. Alternative splicing generates distinct isoforms of the leptin receptor, including long (Ob-Rb) and short isoforms (Ob-Ra and Ob-Rc-f), with Ob-Rb being thought to transmit the majority of leptin’s biological signals (Friedman and Halaas, 1998). However, in addition to its classical role in the neuroendocrine regulation of food intake, the existence of its receptors in extra-hypothalamic brain regions strongly suggests that leptin affects other biological processes. Indeed, studies indicate that leptin is strongly involved in central nervous system (CNS) (Fernandez-Martos et al., 2012; Zhang et al., 2007) and neurological disorders (Fewlass et al., 2004; Greco et al., 2009a; Greco et al., 2009b), through mechanisms of action involving its four major signal transduction pathways: Janus tyrosine kinase/signal transducer and activator of transcription (JAK/STAT) pathway, extracellular signal-regulated kinase pathway (ERK), phosphatidylinositol 3-kinase (PI3K)/Akt pathway, and mitogen-activated protein kinase (MAPK)/sirtuin 1 (SIRT1) pathway, downstream of Ob-Rb receptors (King et al. 2018; Zhang & Chua, 2018). Leptin is reported to be involved in ALS (Lim et al., 2014); however, our understanding of the underlying biological mechanisms of leptin’s actions in the pathogenesis of ALS is limited although both clinical and epidemiological studies support the concept that altered leptin levels contribute to the pathogenesis of ALS (Ngo et al., 2015). A new epidemiological study has determined that leptin levels are inversely associated with ALS outcome (Nagel et al., 2017): increasing leptin concentrations were associated with longer survival of ALS patients, which highlights the possible link between leptin and the clinical outcome of ALS. Altered peripheral levels of leptin have been recently reported in patients with ALS and frontotemporal dementia (FTD) (Ahmed et al., 2019), which exists on a continuous clinical spectrum with ALS (Chen-Plotkin et al., 2010). Interestingly, new epidemiological data suggest that increased dietary fat intake (high fat diet; HFD), which significantly increases leptin levels, may reduce the risk of developing ALS (Morozova et al., 2008; Okamoto et al., 2007; Veldink et al., 2007). A positive correlation between plasma leptin and body mass index (BMI) was observed in ALS patients (Ngo et al., 2015). Nevertheless, a study in SOD1^G93A^ mice has suggested that leptin reduction or loss is beneficial and slows disease progression (Lim et al., 2014), while other studies have investigated the potential therapeutic impact of HFD consumption, which induces obesity and increases leptin levels, in mutant SOD1 mice (Dupuis et al., 2004; Mattson et al., 2007; Zhao et al., 2012; Zhao et al., 2006) and report beneficial effects in terms of survival and improved motor behavior. Therefore, the information to date is not sufficient to clarify the role of leptin or the possible pathways that could link this adipokine to the pathogenesis of ALS. A greater understanding of leptin signaling in ALS is needed to determine whether or not leptin pathways are causally connected to ALS pathogenesis.

In this context, we examined leptin-related pathways during different ALS stages (onset and end-stage of disease) in TDP-43^A315T^ mice (Wegorzewska et al., 2009), which recapitulate several aspects of the human disease. We examined the status of leptin signaling under the backdrop of CNS pathological TDP-43 levels providing, to our knowledge, the first insights into the association between the pathways that could link alterations in leptin signaling to the pathogenesis of ALS.

## METHODS

### Colony maintenance and mice monitoring

Transgenic (Tg) mice TDP43^A315T^ (Strain No. 010700, Bar Harbor, ME, USA) and wildtype (WT) littermate control mice were used in this study (Wegorzewska et al. 2009). This mouse model of ALS expresses a mutant human TAR DNA binding protein TDP-43 cDNA harboring an N-terminal Flag tag and an A315T amino acid substitution associated with ALS mainly in the CNS (Wegorzewska et al. 2009). To avoid ambiguity associated with reported sex-related differences in mean survival time of TDP-43^A315T^ mice (Wegorzewska et al. 2009; Hatzipetros et al. 2014), only male mice were used. Animals expressing the hTDP-43 transgene were confirmed via PCR according to the distributor’s protocol. The ALS-like disease was divided into two stages according to time points: onset (defined as the last day of individual peak body weight before a gradual loss occurs) and end-stage of disease (defined as the weight below 80% of the initial weight on each of three consecutive days). Animals were group-housed under standard housing conditions with a 12 h light–dark cycle, and food and water *ad libitum.* To monitor disease progression and onset determination, all mice were weighed and assessed three times per week until the disease onsetstage, after which they were checked daily in the morning until the disease end-stage. All experimental procedures were approved by the Animal Ethics Committee of the National Hospital for Paraplegics (HNP) (Approval No 26/OH 2018) in accordance with the Spanish Guidelines for the Care and Use of Animals for Scientific Purposes.

### Tissue preparation

Food and water *ad libitum* animals were terminally anesthetized with sodium pentobarbitone (140 mg/kg) and transcardially perfused with room temperature 0.01 M phosphate buffered saline (PBS; pH 7.4), in the middle of the light cycle (between 11 AM and 1 PM). Blood was collected and processed as previously described (Rodriguez et al., 2021). Gonadal white adipose tissue (WAT), hypothalamus and lumbar spinal cord from each animal were processed to extract both mRNA and proteins for real time PCR and Western-blot analysis. Samples were immediately frozen on dry ice and stored at −80°C for later analysis.

### RNA isolation and RT-qPCR

Total RNA was extracted following the instructions of the RNeasyPlus Mini kit (Qiagen, Hilden, Germany). Absorbance at 260 was measured using a NanoDrop (ThermoFisher, Waltham, MA; USA) to determine RNA concentrations.

For WAT and spinal cord samples, complementary DNA (cDNA) was synthesized from 1 μg of total RNA as described previously (Fernandez et al., 2009). Relative quantitation of leptin (assay ID: Mm00434759_m1) and Ob-Rb (assay ID: Mm00440181_m1) was performed using 10 ng of reverse transcribed total RNA in TaqMan One-Step real time PCR Master Mix (PE Applied Biosystem). Each sample was run in duplicate and ß-Actin (assay ID: Mm00607939_s1) was used as a control to normalize gene expression. The reactions were run on an ABI PRISM 7900 Fast Sequence Detection System instrument and software (Applied Biosystem) according to the manufacturer’s protocol.

Furthermore, for the hypothalamus samples, cDNA was synthesized from 1 μg of total RNA by using a NZY First-Strand cDNA Synthesis Kit (NZYTech, Lisbon, Portugal). Quantitative real-time PCR was performed by using assay-on-demand kits (Applied Biosystems). NZY qPCR Probe Master Mix (NZYTech) was used for the PCR reaction was used according to the manufacturer’s protocol in a QuantStudio3 Detection System (Applied Biosystems). Each sample was run in duplicate and glyceraldehyde 3-phosphate dehydrogenase (GAPDH; assay ID: 4352339E) was used as a control to normalize gene expression. The primers and probes used were Ob-Rb (assay ID: Mm00440181_m1), pro-opiomelanocortin (POMC; assay ID: Mm00435874_m1), Agouti-related protein (Agrp; assay ID: Mm00475829_g1), and neuropeptide Y (NPY; assay ID: Mm03048253_m1). In all cases relative quantification for each gene was performed by the ΔΔCt method (Livak and Schmittgen, 2001).

### Protein extraction and western blot analysis

Proteins from spinal cord tissue were extracted with RIPA buffer (Sigma Aldrich) containing a cocktail of protease inhibitors (Roche). For the hypothalamus samples, the supernatant collected in the RNA extraction was diluted in acetone and frozen. Afterwards, samples were centrifuged, and proteins from the hypothalamus were re-suspended in 100 μl of CHAPS buffer (7M urea, 2M thiourea, 4% CHAPS, and 0.5% Tris-HCl 1M, pH 8.8). In all cases, protein concentration was measured using the BioRad Protein assay based on the Bradford method. Samples were measured at 595 nm on an automatic microplate analyzer (Tecan Infinite M200, Grödig, Austria). Denatured protein (30 μg for spinal cord and 20 μg for hypothalamus samples, respectively) were resolved using 8%, 10% or 12% SDS-PAGE (depending on the molecular weight of the protein assayed) and transferred onto PVDF membranes. Membranes were blocked with TBS with 0.1% Tween 20, and 5% BSA or non-fat dried milk and incubated overnight at 4°C with the primary antibody in blocking buffer. The primary antibodies used for spinal cord samples were: rabbit anti-Ob-Rb (1:500; Abcam), rabbit anti-Akt (1:1000; Cell signaling), rabbit anti-Akt (Ser473) (1:1000; Cell signaling), rabbit anti-STAT3 (1:500; Santa Cruz), and rabbit anti-STAT3 (Tyr705) (1:1000; Cell signaling). The primary antibodies used for hypothalamus samples were: rabbit anti-LepR: (1:250, Santa Cruz); Mouse phospho-Akt Ser473 (1:500; Cell Signaling), rabbit anti-Akt (1:1000 Santa Cruz); mouse anti-STAT3 (1:1000; Cell Signaling), and rabbit anti-phospho STAT3 (Tyr705) (1:500; Cell signaling); Rabbit anti-Suppressor of cytokine signaling 3 (SOCS3) (1:1000; Proteintech, Rosemont, IL). A corresponding anti-rabbit or anti-mouse horseradish peroxidase (HRP)-conjugated secondary antibody at a dilution of 1:7000 (Dako) or 1:2000 (Pierce, Rockford II, USA), for spinal cord and hypothalamus samples, respectively, were used. Mouse anti-GAPDH (1:5000, Millipore) and/or mouse anti-actin (1:1000; ThermoFisher) were used as the loading control. Peroxidase activity was visualized by using Immune-Clarity Western Chemioluminiscent substrate (BioRad) and determined by densitometry using an ImageQuant LAS 4000 mini system (GE Healthcare Little Chalfont, United Kingdom). Band intensity was measured as the integrated intensity using ImageJ software (v1.4; NIH). All data were normalized to control values on each membrane.

### Measurement of metabolic markers in plasma

Total ghrelin, the adipokines resistin and leptin, and metabolic biomarkers of insulin resistance (GIP, GLP-1, glucagon, PAI-1 and insulin) from plasma samples were analyzed by duplicate using the Bio-PlexPro mouse Diabetes group from Bio-Rad (Ref. 171F7001M) by Luminex® 200^TM^ technology as previously described (Ortega Moreno et al., 2020). Samples were processed following the manufacturer’s instructions. According to Bio-Rad’s information the intra- and inter-assay CV variability is < 20%. The final concentration value of each metabolic marker was the result of the mean from the two duplicated measures.

### Statistical analysis

All data are presented as means ± standard error of the mean (SEM). Differences between means were assessed by two-way ANOVA followed by Dunett’s post hoc test, to compare all groups with control WT onset mice, and Tukey’s post hoc test were used for multiple comparisons between all groups. For multiplex assays, the mean of each experimental group was determined for all the analytes, and Kruskal–Wallis test was performed followed by Dunett’s post hoc test to compare all groups with onset stage, while Bonferroni post hoc test were used for multiple comparisons between all groups. For all statistical tests, a *p* value of < 0.05 (CI 95%) was assumed to be significant. Statistical analysis was performed using GraphPad Prism software (version 8.3.1).

## RESULTS

### Leptin levels are altered in WAT of TDP-43^A315T^ mice

Experimental data are limited regarding the status of leptin during the progression of ALS, even though this hormone is historically known for its important role in regulating body weight, and mild obesity appears to improve survival in ALS patients (Paganoni et al., 2011). Thus, to determine the leptin expression profile in WAT, the primary source of leptin production, we first examined the expression levels of *leptin* mRNA in different phases of the disease in TDP-43^A315T^ mice compared to age-matched WT littermates (Figure 1). RT-qPCR analysis demonstrated marked differences in the expression profile of the *leptin* transcript during the clinical course of disease in TDP-43^A315T^ mice when compared with WT samples. There was a significant effect of genotype (*p* = 0.001) and disease progression (*p* = 0.002) in the expression profile of *leptin* mRNA in WAT across groups (Figure 1). Indeed, although we (Rodriguez et al., 2021) and others (Esmaeili et al., 2013; Guo et al., 2012; Hatzipetros et al., 2014; Medina et al., 2014) have previously reported that TDP-43^A315T^ mice exhibit weight loss during disease progression. Tukey’s post hoc test demonstrated a statistically significant up-regulation of *leptin* levels in WAT at the onset stage in TDP-43^A315T^ mice, followed by a significant increase in its mRNA expression at the end-stage of disease compared to age-matched WT littermates (*p* = 0.02 and *p* = 0.03, respectively; Figure 1).

**Figure 1.**
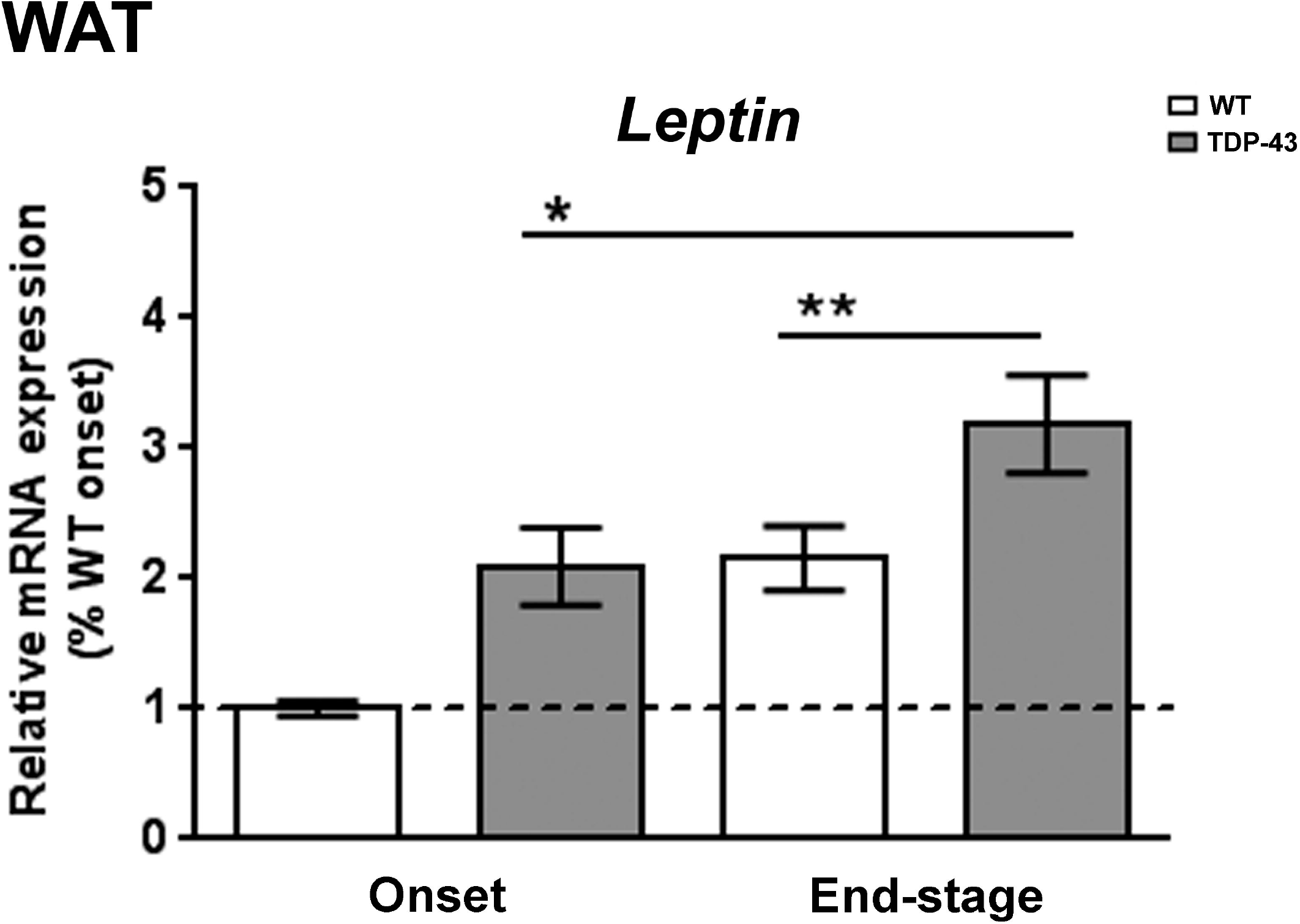
Alterations in *leptin* expression in the WAT in TDP-43^A315T^ mice. *Leptin* mRNA expression was assessed by qRT-PCR in TDP-43^A315T^ mice compared to age-matched WT littermates at both onset and end-stage of disease. Values are expressed as mean ± SEM for the different groups. Comparison between groups was performed by two-way ANOVA followed by Dunett’s post hoc test to compare all groups with WT onset stage, while Tukey’s post hoc test were used for multiple comparisons between all groups, where \**p* < 0.05 vs. WT onset stage; #*p* < 0.05 vs. TDP-43^A315T^ onset; \*\**p* < 0.05 vs. WT end-staged. Abbreviations: WAT, white adipose tissue.

### Peripheral levels of leptin, ghrelin and resistin are altered in plasma of TDP-43^A315T^ mice

Circulating leptin levels were reduced during both ALS stages in TDP-43^A315T^ mice compared to age-matched controls, being significantly lower at disease-termination in the affected mice (F_(3, 12)_=3.875, *p* = 0.03; Figure 2A). Ghrelin and resistin levels also showed differences between the ALS stages, as well as genotype-specific differences (Figure 2B-C). Dunnett’s post hoc test demonstrated a significant increase in total ghrelin concentrations in both WT and TDP-43^A315T^ mice at the end-stage of disease compared to the onset stage (*p* = 0.01 and *p* = 0.007, respectively; Figure 2B). Circulating resistin concentrations were lower in TDP-43^A315T^ mice compared to age-matched WT littermates, with this reaching statistical significance at the end-stage of disease (*p* = 0.001; Figure 2C).

**Figure 2.**
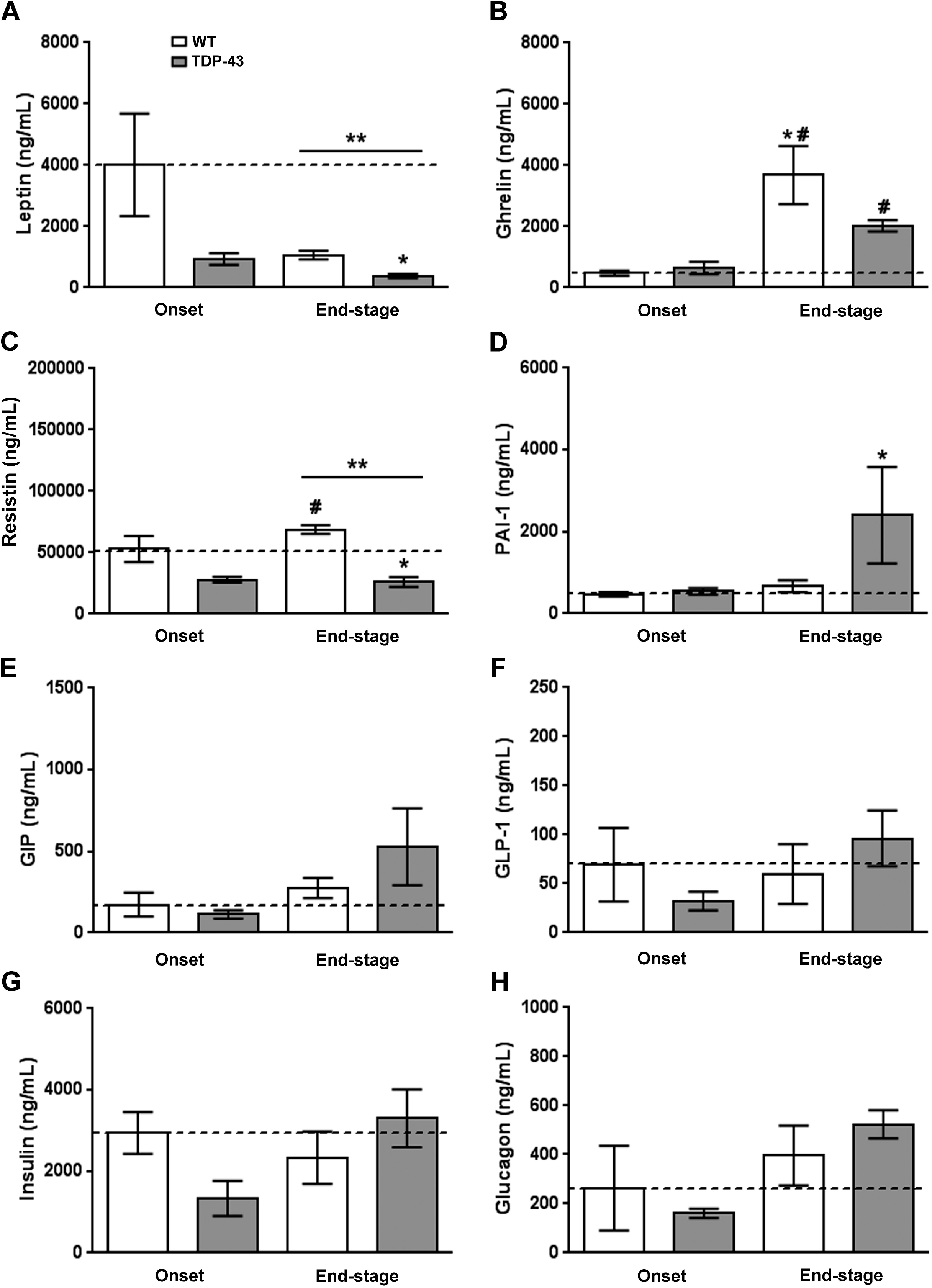
Adipocytokines and metabolic biomarkers levels in WT controls and TDP-43A315T mice. Plasma total ghrelin, the adipokines resistin and leptin, and metabolic biomarkers of insulin resistance (PAI-1, GIP, GLP-1, insulin and glucagon) was measured over the time course of the disease in TDP-43^A315T^ mice compared to age-matched WT littermates using Luminex® 200^TM^ technology. Values are expressed as mean ± SEM for the different groups. Kruskal–Wallis test was performed followed by Dunett’s post hoc test to compare all groups with WT onset stage, while Bonferroni post hoc test were used for multiple comparisons between all groups, where * *p* < 0.05 vs. WT onset stage; #*p* <0.05 vs. TDP-43^A315T^ onset; \*\**p* < 0.05 vs. WT end-staged. Abbreviations: PAI-1, plasminogen activator inhibitor type 1; GIP, gastric inhibitory peptide; GLP-1, glucagon like peptide 1.

To further analyze metabolism, circulating levels of PAI-1, GIP, GLP-1, insulin and glucagon peptides were measured in TDP-43^A315T^ compared to age-matched WT littermates at both time-points of the disease. No statistically significant differences were found between TDP-43^A315T^ and WT mice at either of the time-points analyzed (Figure 2F-H); however, Dunett’s post hoc test demonstrated a significant increase in PAI-1 peptide concentrations in TDP-43^A315T^ mice at the endstage of disease compared to circulating PAI-1 levels in WT mice at the onset stage (*p* = 0.02; Figure 2D). Indeed, no linear correlation was found using Spearman’s test among the plasmatic levels of these metabolic proteins in WT controls or TDP-43^A315T^ mice along the clinical course of disease (data not shown).

### Hypothalamic leptin signaling in TDP-43^A315T^ mice

We next studied how leptin signaling and leptin sensitive genes involved in metabolism were affected in the hypothalamus of TDP-43^A315T^ mice over the time course of the disease. RT-qPCR analysis demonstrated a significant effect of genotype (*p* = 0.001) and disease progression (*p* = 0.002) on the expression profile of *Ob-Rb* mRNA in the hypothalamus (Figure 3A). Dunett’s post hoc test showed that *Ob-Rb* mRNA levels were upregulated in the hypothalamus of TDP-43^A315T^ mice at the both time-points of the disease compared to age-matched WT controls (*p* = 0.03 and *p* = 0.001, respectively; Figure 3A). In addition, as central hypothalamic leptin signaling has a critical role in promoting energy homeostasis via modulation of food intake and energy expenditure (Munzberg et al., 2020), we also investigated the mRNA expression levels of *POMC, AgRP* and *NPY* neuropeptides (Figure 3B-D), that play essential roles in the regulation of food intake and energy homeostasis in mammals. RT-qPCR analysis demonstrated differences in the pattern of expression of *POMC, NPY and AgRP* genes between TDP-43^A315T^ and WT mice at both stages of disease. There were a significant effect of genotype (*p* = 0.03) and disease progression (*p* = 0.0007) in the expression profile of *POMC* mRNA in the hypothalamus across groups (Figure 3B), with an overall increase in *POMC* mRNA with age in both genotypes. In TDP-43^A315T^ mice *POMC* mRNA levels were lower than in WT at both onset and end-stage, with this increase being significant at the end-stage of disease compared to *POMC* mRNA levels in WT controls at the onset (*p* = 0.04; Figure 3B). In addition, RT-qPCR analysis demonstrated a significant effect of genotype (*p* = 0.0009 and *p* = 0.0005, respectively) and disease progression (*p* = 0.0004 and *p* = 0.002, respectively; Figure 3C and D) on the expression profile of *NPY* and *AgRP* mRNA levels in the hypothalamus. Although Dunett’s post hoc test demonstrated no age dependent changes in *NPY* and *AgRP* mRNA levels in TDP-43^A315T^ mice compared to WT mice at the onset stage, Tukey’s post hoc test demonstrated a statistically significant hypothalamic upregulacion of both orexigenic neuropeptides (*NPY: p* = 0.002, Figure 3C; and *AgRP: p* = 0.0007, Figure 3D) at the end-stage of disease in the hypothalamus of TDP-43^A315T^ mice relative to age-matched WT controls.

**Figure 3.**
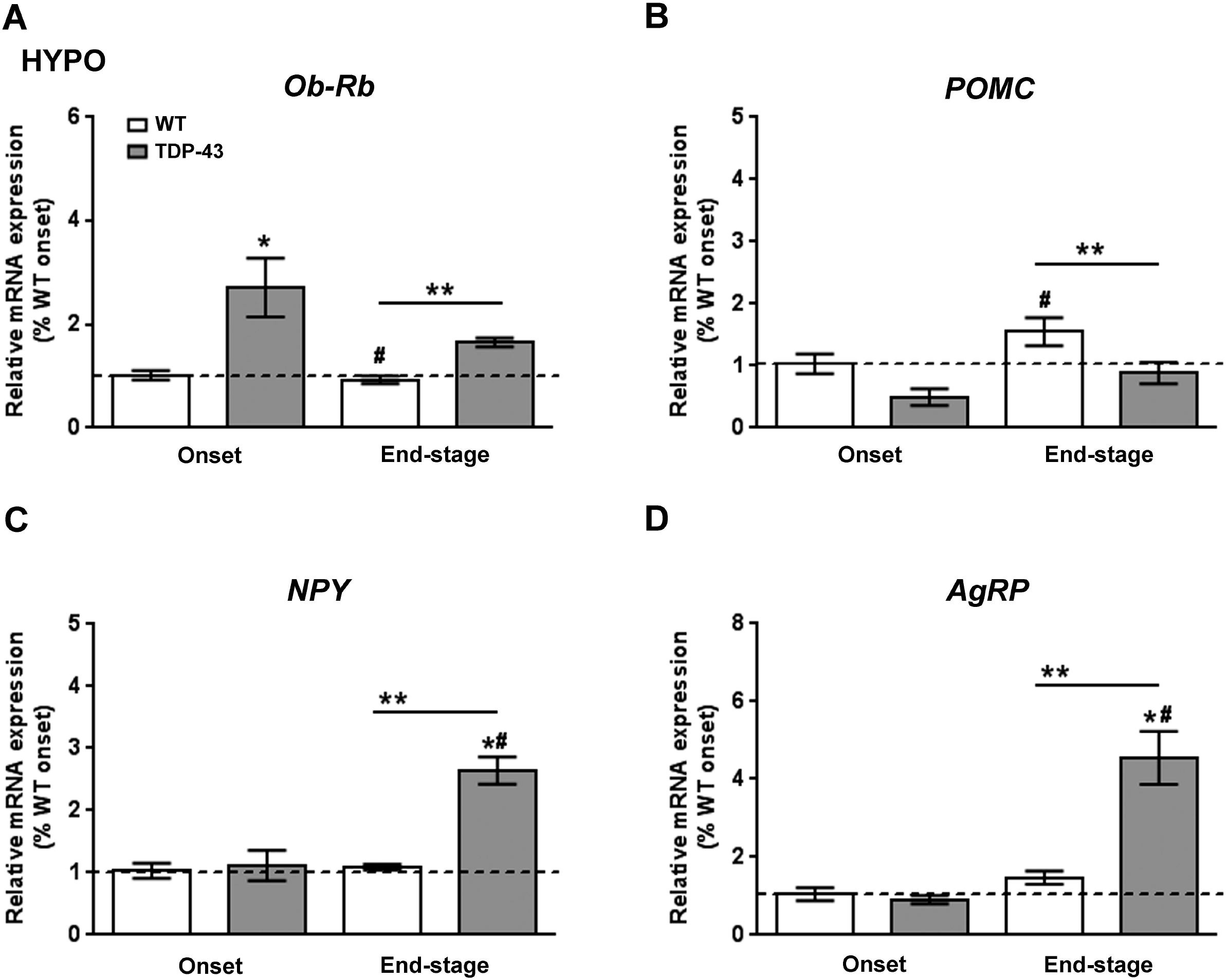
Alterations in Ob-Rb and anorexigenic and orexigenic neuropeptides in the hypothalamus of TDP-43^A315T^ mice. (A) *Ob-Rb* mRNA expression, (B) *POMC,* (C) *NPY* and (D) *AgRP* transcripts were assessed by qRT-PCR in TDP-43^A315T^ mice compared to age-matched WT littermates at both onset and end-stage of disease. Values are expressed as mean ± SEM for the different groups. Comparison between groups was performed by two-way ANOVA followed by Dunett’s post hoc test to compare all groups with WT onset stage, while Tukey’s post hoc test were used for multiple comparisons between all groups, where \**p* < 0.05 vs. WT onset stage; *p* <0.05 vs. TDP-43^A315T^ onset; \*\**p* < 0.05 vs. WT end-staged. Abbreviations: HYPO, hypothalamus; Ob-Rb, long form of leptin receptor; POMC, Proopiomelanocortin; Agrp, Agouti-related protein; NPY, Neuropeptide Y.

We next investigated the protein levels of Ob-Rb (Figure 4A), SOCS3 (Figure 4B), a main inhibitor of leptin signaling in the brain, as well as, the status of the STAT3 (pTyr^705^-STAT3) and Akt (pSer^473^-Akt) pathways, which are downstream of the Ob-Rb receptor (Figure 4C-D). Immunoblotting analysis demonstrated that Ob-Rb was also altered at the protein level in TDP-43^A315T^ mice (Figure 4A). There was a significant effect of disease progression (*p* = 0.005) on the expression profile of Ob-Rb receptor in the hypothalamus. In contrast to mRNA levels, protein levels of Ob-Rb receptor were lower in TDP-43^A315T^ mice compared to age-matched WT controls, which reached significance (*p* = 0.01) at the end-stage of disease (Figure 4A). In addition, we found no significant effect of either disease stage or genotype on SOC3 levels (Figure 4B). There was a significant effect of genotype (*p* = 0.0001) on the phosphorylation levels of Akt protein in the hypothalamus (Figure 4C). At the end-stage of disease, Akt phosphorylation levels were significantly decreased in the hypothalamus of both genotypes (*p* = 0.001 and *p* < 0.0001, respectively) compared to WT controls at the onset stage (Figure 4C). Furthermore, Akt phosphorylation was also significantly decreased when comparing TDP-43^A315T^ mice at different time-points (*p* = 0.0008), showing an effect of disease course on this pathway (Figure 4C). In addition, there was no effect of genotype and disease progression in the expression of phosphorylation levels of STAT3 protein in the hypothalamus across groups (Figure 4D).

**Figure 4.**
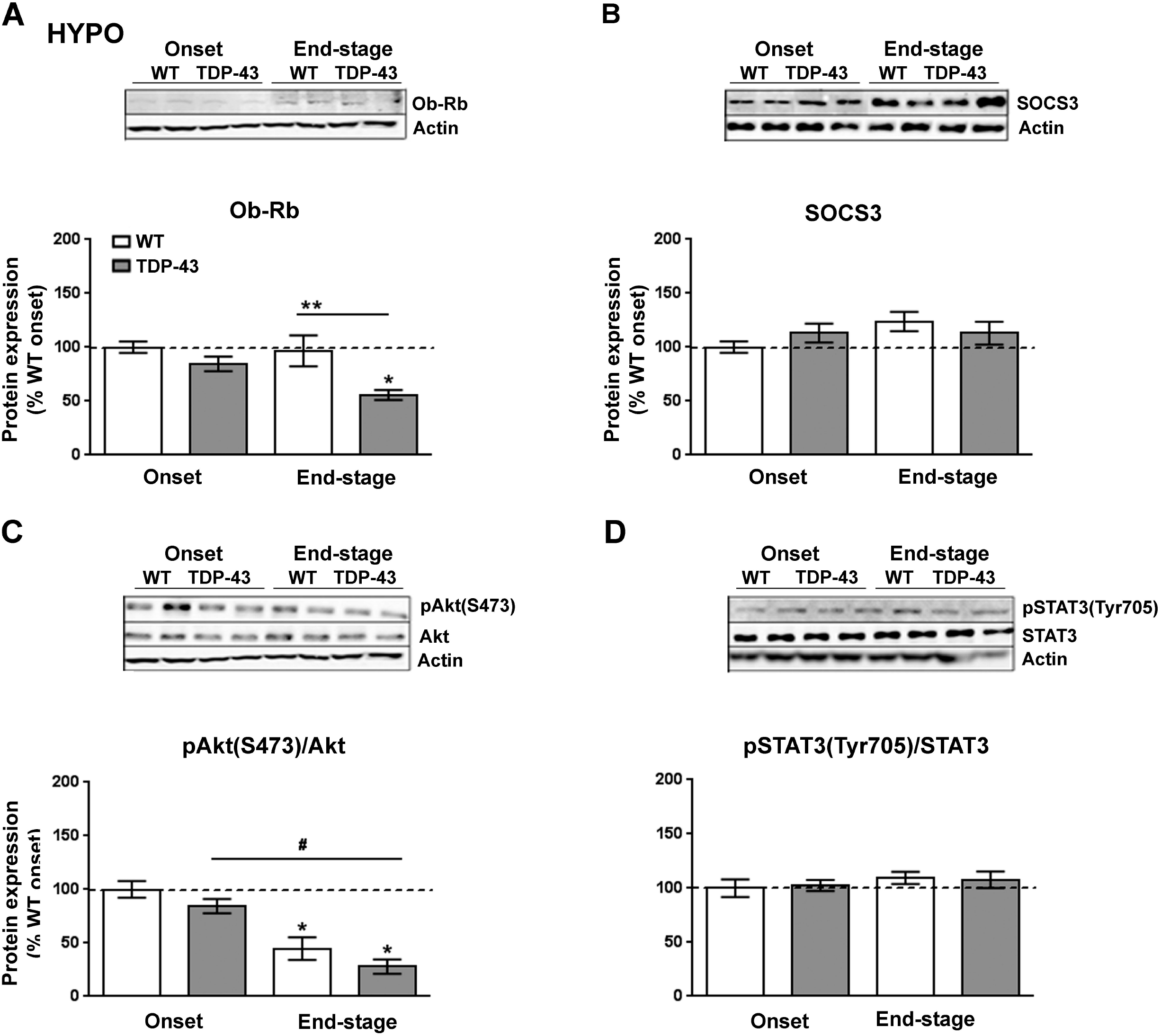
Alterations in serine phosphorylation of Akt in the hypothalamus of TDP-43^A315T^ mice. Representative Actin-normalized immunoblot images and quantitation of (A) Ob-Rb receptor, (B) SOCS3, (C) pAkt (pSer^473^-Akt) protein, (D) pSTAT3 (pTyr705-STAT3) proteins, respectively, in hypothalamic extracts of TDP-43^A315T^ mice compared to age-matched WT littermates at both onset and end-stage of disease. Values are expressed as mean ± SEM for the different groups. Comparison between groups was performed by two-way ANOVA followed by Dunett’s post hoc test to compare all groups with WT onset stage, while Tukey’s post hoc test were used for multiple comparisons between all groups, where \**p* < 0.05 vs. WT onset stage; #*p* <0.05 vs. TDP-43^A315T^ onset; \*\**p* < 0.05 vs. WT end-staged. In the immunoblot images, representative bands were run on the same gel but were non-contiguous. Abbreviations: HYPO, hypothalamus; Ob-Rb, long form of leptin receptor; SOCS3, Suppressor of cytokine signaling 3; Akt, Serine*/*threonine kinase; STAT3, Signal transducer and Activator of transcription 3.

### Leptin signaling in the spinal cord of TDP-43^A315T^

Since leptin signaling has actions throughout the CNS (Zhou and Rui, 2013), and the results of our study could possibly indicated that the reduction in circulating leptin levels are associated with altered hypothalamic leptin signaling in TDP-43^A315T^ mice, particularly at the end-stage of disease, we analyzed if leptin signaling in the spinal cord tissue of TDP-43^A315T^ mice differed from that of WT mice. Although there was no effect of genotype, RT-qPCR analysis demonstrated that there was a significant effect of disease progression (*p* = 0.01) on the expression profile of Ob-Rb receptor in the spinal cord (Figure 5A). In addition, Tukey’s post hoc test demonstrated a statistically significant down-regulation of *Ob-Rb* mRNA at the end-stage of disease in the spinal cord of TDP-43^A315T^ relative to age-matched WT littermates (*p* = 0.02, Figure 5A). In contrast, Ob-Rb protein levels were increased in the spinal cord of TDP-43^A315T^ mice compared to WT at both onset and end-stage of the disease (*p* = 0.001 and *p* = 0.04, respectively; Figure 5B). In addition, there was a significant effect of genotype (*p* = 0.005) and disease progression (*p* = 0.0006) on the expression of phosphorylation levels of Akt protein in the spinal cord across groups (Figure 5C). Finally, there was a significant effect of genotype (*p* = 0.0001; Figure 5D) on the expression of phosphorylation levels of STAT3 protein in the spinal cord across groups, with levels increasing in both genotypes with age. Indeed, Dunnett’s post hoc test demonstrated that phosphorylation levels of STAT3 protein was significantly decreased at the end-stage of disease in TDP-43^A315T^ compared to WT controls at the onset stage (*p* = 0.004; Figure 5D).

**Figure 5.**
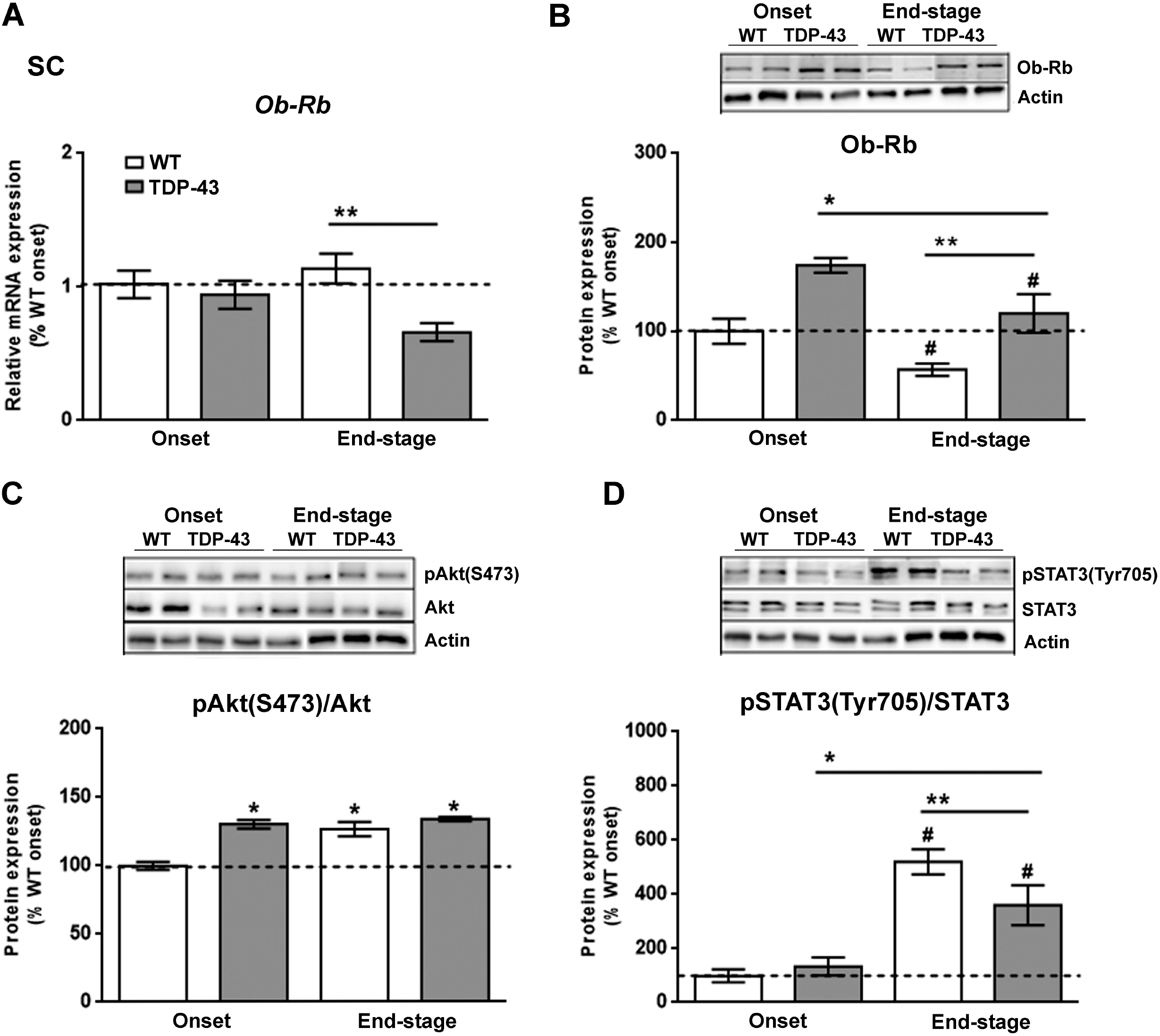
Alterations in tyrosine phosphorylation of STAT3 in the spinal cord of TDP-43^A315T^ mice. (A) mRNA expression of *Ob-Rb* receptor was assessed by qRT-PCR in TDP-43^A315T^ mice compared to age-matched WT littermates at both onset and end-stage of disease. Representative GAPDH-normalized immunoblot images and quantitation of (B) Ob-Rb receptor, (C) pAkt (pSer^473^-Akt) protein, (D) pSTAT3 (pTyr705-STAT3) protein in spinal extracts of TDP-43^A315T^ mice compared to age-matched WT littermates at the onset and end-stage of disease. Values are expressed as mean ± SEM for the different groups. Comparison between groups was performed by two-way ANOVA followed by Dunett’s post hoc test to compare all groups with WT onset stage, while Tukey’s post hoc test were used for multiple comparisons between all groups, where *p* < 0.05 vs. WT onset stage; #*p* < 0.05 vs. TDP-43^A315T^ onset; \*\**p* < 0.05 vs. WT end-staged. In the immunoblot images, representative bands were run on the same gel but were non-contiguous. Abbreviations: SC, spinal cord; Ob-Rb, long form of leptin receptor; SOCS3, Suppressor of cytokine signaling 3; Akt, Serine/threonine kinase; STAT3, Signal transducer and Activator of transcription 3.

## DISCUSSION

A growing body of evidence shows disturbances in energy metabolism in ALS (Blasco et al., 2020; Tefera and Borges, 2016; Tefera et al., 2021; Vandoorne et al., 2018), suggesting that targeting metabolism could represent a rational strategy to treat this disease. Metabolic abnormalities have been reported in both ALS patients (Dupuis et al., 2004) and mouse models of ALS (Lim et al., 2014), as well as in the more recently developed murine model of ALS/FTD, TDP-43 proteinopathy (Shan et al., 2010; Wang et al., 2013). Two epidemiological studies provided the first evidence of leptin as a potential novel therapeutic target in ALS (Bejanin et al., 2020; Nagel et al., 2017), although very little is known about the direct influence of leptin in altering energy metabolism and disease progression in ALS, as it has thus far been correlated with the protection exerted by increased fat mass stores. Indeed, even though leptin signaling appears to be involved in ALS, our understanding of its biological role in mechanisms of disease pathogenesis is limited. Here, we present evidence of alterations in leptin signaling in the peripheral and CNS of the TDP43^A315T^ transgenic ALS mouse model, providing novel insights about the pathways that could link alterations in leptin to ALS disease.

The majority of circulating leptin is produced in adipose tissue (Zhang and Chua, 2017), with *leptin* mRNA levels normally being directly correlated with adipocyte size, and high circulating levels of this hormone are associated with obesity (Cohen et al., 2017; Zhang and Chua, 2017). The opposite is observed in ALS patients (Lopez-Gomez et al., 2021; Ludolph et al., 2020) as ALS causes loss of body weight, reduced fat mass, and reduced circulating leptin levels. Here we report an up-regulation of *leptin* mRNA levels in WAT of TDP-43^A315T^ mice, both at onset and at the end-stage of the disease. This observation is of interest because peripheral leptin levels are positively correlated with adipose tissue mass in TDP-43^A315T^ mice, as we previously reported a progressive decline in body weight in TDP-43^A315T^ mice compared to WT controls (Rodriguez et al., 2021). Indeed, circulating plasma levels of leptin were lower in TDP-43^A315T^ mice compared to WT mice at both ALS stages, which is in accordance with the decrease in body weight (Esmaeili et al., 2013; Guo et al., 2012; Hatzipetros et al., 2014; Medina et al., 2014). Nevertheless, although the reduction in circulating levels of leptin is in accordance with a lower adipose tissue mass, the mRNA levels of leptin/μg adipose tissue is upregulated in TDP-43^A315T^ mice that might suggest an attempt to maintain normal circulating levels of this adipokine. This observation is of interest because evidence supports that cachexia may occur in the early course of ALS (Holm et al., 2013), even before the loss of motor neurons and neurodegeneration (Ferri and Coccurello, 2017). Indeed, WAT is specialized in the storage of triglycerides (TGs) (Pundir and Narwal, 2018) and patients with ALS suffer from hypolipidemia (Vandoorne et al., 2018). In this context, as adipose tissue wasting has been shown to occur before the appearance of classical cachexia markers as for example loss of fat mass, and subsequently, loss of body weight, it will be interesting in future *in vitro* studies to determine the mechanism of leptin regulation in primary adipocytes of mutant TDP-43. Considering that cachexia may occur in the early course of ALS, our results might suggesting leptin as a potential biomarker of adipose tissue wasting, and subsequently, the muscle atrophy and depletion of fat stores clinical features in ALS.

In addition to the marked decrease of circulating leptin concentrations, our results confirm disease stage-dependent alterations in the circulating levels of ghrelin and resistin in TDP-43^A315T^ mice. Plasma levels of ghrelin, an appetite stimulating hormone, were highest in WT animals at end-stage of disease and although there was an increase between onset and end-stage in TDP-43^A315T^ mice they remained significantly lower compared to WT, which could partly due to modifications in food intake and ultimately the loss of body weight in TDP-43^A315T^ mice (Esmaeili et al., 2013; Guo et al., 2012; Hatzipetros et al., 2014; Medina et al., 2014). Indeed, low plasma ghrelin levels have been found in ALS patients (Ngo et al., 2015). We also found lower circulating levels of the adipokine resistin in TDP-43^A315T^ mice both at onset and end-stage of ALS disease. This result support previous data from our group showing a downregulation of peripheral protein resistin levels in TDP-43^A315T^ mice (Rodriguez et al., 2021). However, although no difference in plasmatic levels of resistin was found between controls and ALS patients (Ngo et al., 2015), this data might indicate that resistin levels are directly associated with metabolic abnormalities in TDP-43^A315T^ mice. However, future experiments should try to corroborate this hypothesis.

We examined leptin signaling in the hypothalamus and spinal cord of TDP-43^A315T^ mice, as they represent two areas of the nervous system vulnerable to ALS disease, in which leptin could play an important role. In both tissues, there was an up-regulation in the expression levels of *Ob-Rb* transcript in TDP-43^A315T^ mice compared to age-matched WT littermates. The observation in spinal cord is of particular interest, as a previous study conducted in rats showed a significant upregulation of *Ob-Rb* mRNA after spinal cord injury (Fernandez-Martos et al., 2012) and thus, our result may reflect the progressive irreversible neurodegenerative damage that characterizes ALS pathogenesis. Alternative splicing of the *ObR gene* generates distinct isoforms of the leptin receptor, including long (Ob-Rb) and short isoforms (Ob-Ra and Ob-Rc-f) that differ in the length of their intracellular cytoplasmic domains, a region that contains specific motifs involved in leptin signaling (Lee et al., 1996; Tartaglia et al., 1995). In this context, the increase in *Ob-Rb* mRNA expression, but decrease in protein levels, could indicate an increase in Ob-Rb receptor turn-over or a modification in processing -producing less long-form and more of the other forms. However, future experiments are necessary to corroborate this hypothesis.

Alternatively, although the precise dynamics of Ob-Rb regulation in both areas of the nervous system are not completely understood, TDP-43 pathology and consequently the progression of ALS stages may be related to leptin signaling disruption. Of particular interested are the Akt and STAT3 pathways downstream of the Ob-Rb receptor, as they are important targets in the regulation of glucose and energy metabolism (Varela and Horvath, 2012). Indeed, we have previously reported that TDP-43^A315T^ mice are hypoglycemic compared to WT mice at the disease end-stage, confirming the disturbances in energy metabolism of the TDP-43^A315T^ mouse model reported previously (Chiang et al., 2010). Here, a significant decrease in serine phosphorylation of Akt was found in TDP-43^A315T^ mice at end-stage, while no differences were founded between genotypes in the spinal cord tissue over the time course of disease, which could represent the different physiological roles that leptin exerts in these two brain areas. In addition, while no differences in hypothalamic tyrosine phosphorylation of STAT3 were observed over the time course of the disease, an increased phosphorylation of STAT3 was observed in the spinal cord of TDP-43^A315T^ and WT mice compared to the onset stage, as well as at the end-stage of disease, in accordance with previous research conducted in SOD1^G93A^ mice (Ohgomori et al., 2018; Ohgomori et al., 2017).

It is conceivable that changes in leptin signaling in the spinal cord of TDP-43^A315T^ mice could potentially to be due to a direct effect of leptin on alpha motor neurons. Indeed, we are currently evaluating the presence of Ob-Rb protein in TDP-43^A315T^ mice by immunohistochemical stainings, and our preliminary unpublished data indicate the presence Ob-Rb receptor in a minority of cells in the ventral horn of the spinal cord tissue that have morphological characteristics of alpha motor neurons in TDP-43^A315T^ and WT mice. In addition, Ob-Rb immunoreactive cells appear to be glial, either a subpopulation of astrocytes or microglia in both genotypes (data not shown). However, further immunohistochemistry studies investigating Ob-Rb localization within spinal cord between genotypes, may be warranted.

Leptin is reported to activate POMC neurons and inhibit both AgRP/NPY neurons, at least in part through the STAT3 pathway (Varela and Horvath, 2012). Indeed, the regulation of glucose homeostasis is related to leptin signaling in the hypothalamic POMC, NPY and AgRP neurons. Neuropeptides derived from POMC provide a strong anorexigenic effect (i.e., decreases food intake), while NPY and AgRP neurons have a potent orexigenic effect (i.e., increase food intake). Thus, in a situation of negative energy balance, such as the malnutrition observed in ALS, the expression of NPY and AgRP is normally increased and POMC expression decreased (Caron et al., 2018; Pedroso et al., 2016). Consistently, our data showed an up-regulation of *NPY* and *AgRP* in TDP-43^A315T^ mice at disease end-stage, as previously reported by others in ALS patients and several animal models (Clark et al., 2021; Vercruysse et al., 2016) and an overall decrease in *POMC* expression. These alterations in metabolic neuropeptides could partly explain the hypoglycemic state observed in TDP-43^A315T^ mice (Rodriguez et al., 2021). Collectively, these transcriptional modifications, at the end-stage of disease in TDP-43^A315T^ mice, perhaps reflect the physiological response of the hypothalamus to overcome from adipose atrophy and loss of body weight.

## CONCLUSSIONS

In summary, our study provides the first experimental evidence suggesting that ALS may be associated with alterations in leptin signaling pathways that might result in a leptin resistant state and that this could play a critical role in the irreversible and progressive characteristic pathological changes associated with this disease. However, the precise pathways that could link leptin signaling to the TDP-43 proteinopathy model of ALS remain unclear. Further mechanistic studies analyzing the consequences of leptin signaling alterations may require at the inclusion of additional defined time-points, as well as larger sample sizes. Determining the role of leptin and its mechanistic actions may provide a new avenue for therapeutic development for this fatal condition.

## Acknowledgments

The authors would like to gratefully acknowledge Sandra Canelles (CIBEROBN, Hospital Infantil Universitario Niño Jesús) and the Animal Facility and Experimental Surgery Unit of the UDI-HNP for their excellent technical support.

## Author contribution

Conceptualization: CM.F.-M. Data curation: CM.F.-M., A.F.-D. Formal analysis: A.C., L.M.F, CM.F.-M. Funding acquisition: J.A.C., CM.F.-M. Investigation: A.C., L.M.F., CM.F.-M. Methodology: A.F.-D., A.C., L.M.F, CM.F.-M. Project administration: CM.F.-M. Resources: J.A.C., CM.F.-M; Software: A.C., L.F.M.; Supervision: J.A.C., CM.F.-M. Validation: A.C., L.M.F., J.A.C., CM.F.-M. Visualization: A.C., L.M.F., J.A.C., CM.F.-M. Writing - original draft: A.C., CM.F.-M. Writing - review & editing: A.C., L.M.F., J.A.C., CM.F.-M.

